# A high-throughput fluorescent indicator displacement assay and principal component/cluster data analysis for determination of ligand-nucleic acid structural selectivity

**DOI:** 10.1101/192849

**Authors:** Rafael del Villar-Guerra, Robert D. Gray, John O. Trent, Jonathan B. Chaires

## Abstract

We describe a high-throughput fluorescence indicator displacement assay (HT-FID) to evaluate the affinity and the selectivity of compounds binding to different DNA structures. We validated the assay using a library of 30 well-known nucleic acid binders containing a variety chemical scaffolds. We used a combination of principal component analysis and hierarchical clustering analysis to interpret the results obtained. This analysis classified compounds based on selectivity for AT-rich, GC-rich and G4 structures. We used the FID assay as a secondary screen to test the binding selectivity of an additional 20 compounds selected from the NCI diversity set III library that were identified as G4 binders using a thermal shift assay. The results showed G4 binding selectivity for only a few of the 20 compounds. Overall, we show that this HT-FID assay, coupled with PCA and HCA, provides a useful tool for the discovery of ligands selective for particular nucleic acid structures.

## INTRODUCTION

Nucleic acids are dynamic biomolecules that can adopt a wide variety of structures (1-3). In addition to the familiar duplex structure, nucleic acids can form a variety of non-canonical triplex (4,5), quadruplex (6) or i-motif (3,7,8) structures. Some of these structures have been detected in living cells (9-11) where they may play key roles in a variety of biological processes, including replication, transcription, oncogene expression, and telomere functions (11-17). Identification of ligands that selectively bind to duplex, triplex, i-motif and G-quadruplex (G4) structures is an important and challenging goal of drug discovery (12,18-32). G4 structures, in particular, have emerged as promising therapeutic targets for anti-cancer therapies (13,18,27-29,33-42). A major challenge is to identify compounds that bind specifically to one of the several possible G4 structures without binding to duplex DNA. Over the last decades, a number of ligands that bind G4 structures have been described (38,43). Unfortunately, none have successfully progressed completely through clinical trials to become useful drugs.

This failure might be because most G4 binders found to date are polyaromatic compounds with poor drug-like properties and nonselective binding (42). Indeed, few G4 binders have been rigorously evaluated for selectivity with respect to their interaction with other nucleic acid structures, in part because there is no convenient assay for evaluating their structural-selective binding. A number of potential binding assays have been developed including chip-based (44), ligand fishing (45), FRET melting (46) and equilibrium dialysis (21,23,25). However, these methods may be expensive, time consuming or not amenable to high-throughput screening.

The fluorescent indicator displacement assay (FID) (27,28,47-56), which relies on displacement of a fluorescent DNA-binding ligand by the test compound, was adapted for the discovery of new G4 binding ligands. However, most previous FID studies evaluated binding only to a single G4 structure, but did not assess binding to other DNA structures or to different G4 folds. Here we describe an efficient and inexpensive high-throughput FID assay (HT-FID) designed to explore the structural selectivity of binding. The assay measures the affinity and selectivity of a library of compounds against eight different DNA sequences and structures (4 duplex, 2 triplex and 2 G4 structures). It was tested using a library of 30 compounds with defined DNA binding modes and preferences.

Importantly, we also developed a chemometric approach for evaluation of the large data set to facilitate visualization of the affinity, structural selectivity and binding mode of the compound library. Traditional graphical methods for the analysis of large data sets are often ineffective for easily visualizing binding selectivity. Data analysis therefore becomes a bottleneck in drug discovery. We present here a novel application of principal component analysis (PCA) and hierarchical cluster analysis (HCA) as powerful tools for analysis of large FID data sets. We demonstrate that PCA and HCA can be used to visualize binding properties and to clearly identify the binding selectivity of a library of compounds. This visualization provides a direct and easily interpretable method to depict complex, often hidden, relationships between affinity and selectivity.

One may question why yet another variation of the FID assay needed. As part of a drug discovery platform it is essential, once a ligand is found that hits the desired target, to evaluate its binding specificity. We envision this FID assay as a high-throughput secondary screen for that purpose. Primary high-throughput screening of large compound libraries for target hits can be reliably done by thermal shift assays, particularly differential scanning fluorometry. The FID assay we describe will fill an important need by providing a high-throughput specificity screen to identify the best candidates to continue along the drug discovery platform.

## MATERIAL AND METHODS

### Preparation of oligonucleotides samples

The sequences, names of the oligonucleotides and buffers used in this study are given in Table 1. Oligonucleotides were purchased as desalted lyophilized powders from Integrated DNA Technologies, Inc. (Coralville, IA) and used without further purification. Stock solutions were prepared at 2 mM concentration in Milli-Q water and stored at 4°C for a least 24 h before use. Further dilutions to working concentrations were prepared using the following buffers optimized for specific DNA structures: Buffer A (6 mM K_2_HPO_4_, 4 mM KH_2_PO_4_, 15 mM KCl, pH7. 2), Buffer B (1 mM K_2_HPO_4_, 9 mM KH_2_PO_4_, 15 mM KCl, pH6.2), Buffer C (6 mM K_2_HPO_4_, 4 mM KH_2_PO_4_, 15 mM KCl, 1 mM MgCl_2_, pH=7. 2). To induce quadruplex, triplex or duplex formation, stocks solutions of each oligonucleotide were diluted with the appropriate buffer to ∼200 μM and annealed by heating in a boiling water bath for 20 min followed by overnight cooling to room temperature. The annealed samples were stored at 4°C. DNA strand concentrations were estimated from the absorbance at 260 nm after thermal denaturation for 5 min at 90 °C using molar extinction coefficients supplied by the manufacture. Oligonucleotide solutions that contained 1% (v/v) dimethyl sulfoxide (DMSO) were prepared by adding the necessary volume of DMSO to the previously annealed nucleic acid solution. All nucleic acid structures were characterized by circular dichroism and UV absorbance spectroscopies, and by thermal denaturation (see Supplementary Material).

**Table 1.**
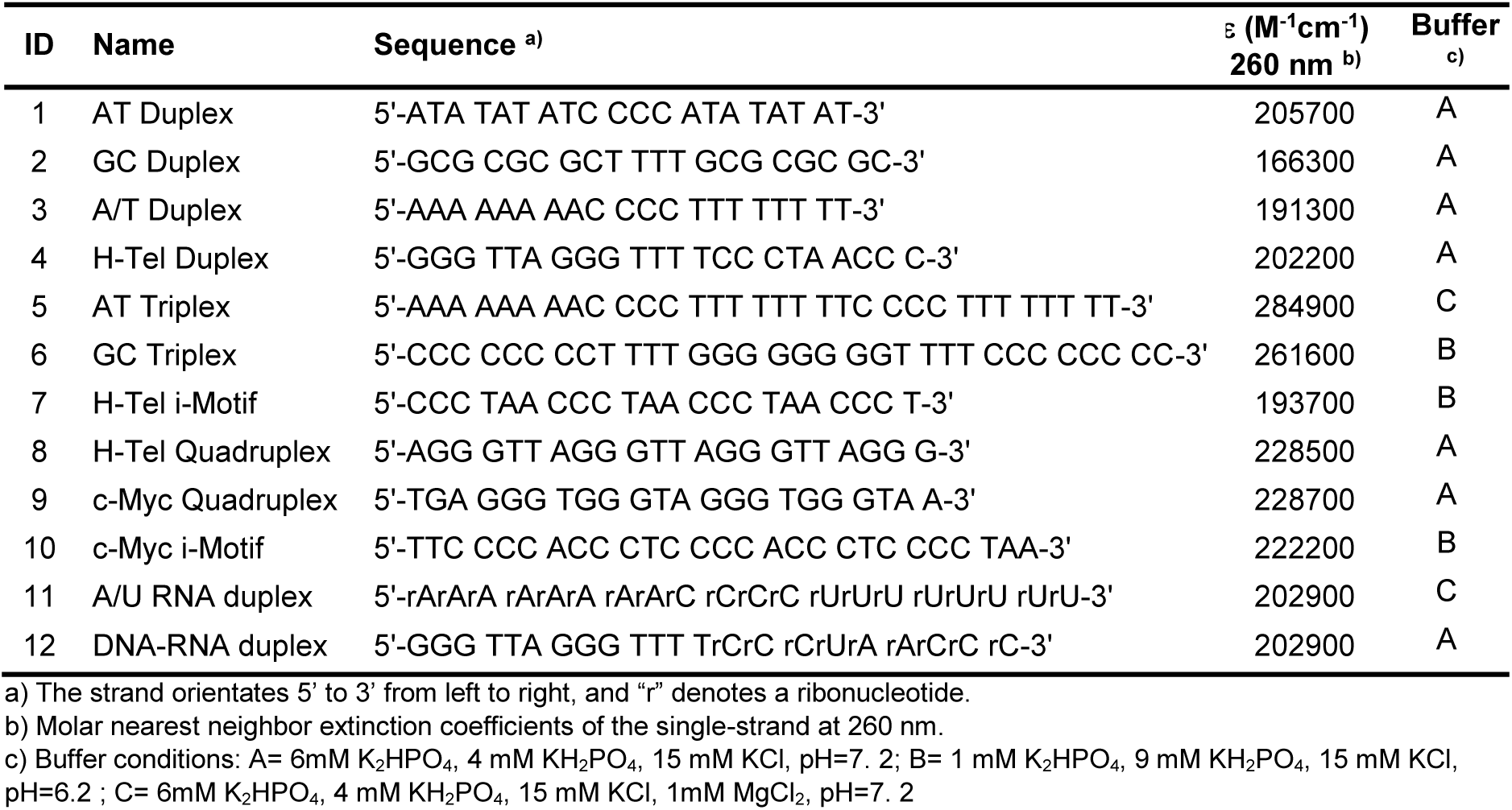
Oligonucleotides sequence and their molar extinction coefficients used in this study

### Sample Preparation

Thiazole Orange (TO) and DMSO (99.9 % purity) were purchased from Sigma-Aldrich (St. Louis, MO) and used without further purification. The test compounds were purchased from commercial sources and were used without further purification. Stock solutions of test compounds were prepared by dissolving a weighed amount in DMSO to give a concentration of 1-10 mM and stored in the dark at -20 °C to prevent light-induced degradation. Working solutions of test compounds were prepared immediately before use at 15 μM concentration in the appropriate buffer containing 1% (v/v) DMSO. Serial dilutions of the stock ligand solutions contained1% (v/v) DMSO. Stock solutions for FID screening were made in the same buffer used for the DNA preparation (Table 1). Oligonucleotide stock solutions (6 μM) were prepared in the appropriate buffer solutions with 1% (v/v) of DMSO using the pre-folded oligonucleotide stock solution (200 μM) prepared as described above.

### Thiazole orange displacement assay

FID assays was conducted in duplicate at room temperature (25-30 °C) with a Tecan Safire^2^ microplate reader (Tecan US, Durham, NC, USA) with the following parameters: excitation and emission bandwidth of 5 nm, gain of 125, integration time of 200 μs, and 16 reads. Emission spectra were measured at 1 nm intervals from 510 to 750 nm with an excitation wavelength of 500 nm. Assays were carried out in 96-well NBS™ black, flat bottom polystyrene microplates (Cat# 3650, Corning Inc., NY, USA). Each assay solution contained 1 μM TO, 2 μM oligonucleotide and 5 μM test compound in 150 μL. The instrumental gain was adjusted using the fluorescence of the most fluorescent sample (1 μM TO and 2 μM AT triplex 5. A representation of a 96-well plate assay is depicted in Figure S1. This configuration allowed testing five compounds and eight oligonucleotide structures per plate along with appropriate controls.

The percentage TO displacement (%FID) was calculated from the corrected fluorescence emission intensity at 527 nm (λ_ex_ = 500 nm) using **Equation 1**. The fluorescence intensity F was corrected by subtracting the fluorescence signal of the test compound/DNA mixture and that of unbound TO:

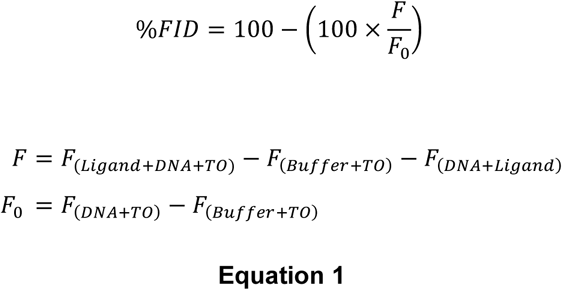

The %FID was calculated using the fluorescence intensity at the TO emission maximum of 527 nm rather than using the integrated intensity over the wavelength range 510-750 nm as has been used in previous implementations of the FID method (28,47,49,54,57). We observed that interference can be introduced when the integrated intensity is used instead of the single wavelength intensity, especially when fluorescent compounds are tested (Figure S2-3).

### Principal component analysis (PCA) and cluster analysis

Multivariate data analysis, principal component analysis, and cluster analysis of the FID assay data were performed with R software version 3.2.3 (2015-12-10), using the R package FactoMineR (58). These programs are freely available from the Comprehensive R Archive Network (CRAN) at http://cran.r-project.org.

Hierarchical clustering on the principal components (59) was performed on the principal components using the Euclidean distance as a measure of similarity between individuals with the Ward criterion as agglomeration method. The initial clusters obtained by hierarchical clustering were further consolidated by partitional clustering using the *k*-means algorithm. The results of the cluster analysis are presented as dendrograms in which the horizontal axis represents the compounds and the vertical axis represents the degree of similarity of the individuals.

## RESULTS AND DISCUSSION

### Oligonucleotide design and characterization

A library of 12 oligonucleotides that form representative duplex, triplex, i-motif and G4 structures was designed for use in the new FID assay. Unimolecular structures were selected to avoid potential problems related to concentration and the molecularity of folding. Hairpin duplex structures are represented by deoxyribooligonucleotides 1-4 and the RNA oligonucleotides 11-12. These structures represent B-form DNA with AT- and GC-rich base compositions, an A-form RNA and a DNA-RNA hybrid. Sequences 5 and 6 form AT and GC intramolecular triplex DNA structures, respectively. These triplex structures (H-DNA structures) are important in several biological processes (60). Oligonucleotides 7 and 10 represent i-motif structures (7,61) corresponding to the human telomeric sequence (62) and the c-MYC NHE III1 wild-type C-rich promoter sequence (3). Oligonucleotides 8 and 9 form representative G4 structures. H-Tel (8) is the human telomeric repeat sequence (63,64) that is important in cancer (16,39) and which is polymorphic depending on environmental conditions (63,65-67) An antiparallel hybrid (“3+1”) structure is formed under the conditions of the assay. c-Myc (9) is a sequence variant of the wild-type NHE III1 sequence of the c-Myc promoter that forms a homogeneous parallel G-quadruplex structure in potassium solution (68).

Each of these nucleic acid structures have different stabilities and require optimization of experimental conditions to ensure their formation. The influence of buffer composition, pH, DMSO and TO on the folding and stability of the structures was evaluated using UV-Vis absorption, circular dichroism spectroscopy (CD) (69-74), thermal difference spectra (TDS) (75), and UV/CD thermal denaturation (Figures S4-15). A summary of the physical and spectroscopic characteristics of the oligonucleotides under different buffer conditions is given in Table S2.

The effect of TO and DMSO (used as a solvent for the ligands) on the stability and folding of the triplex oligonucleotides was also evaluated (Figure S16-18). CD and melting experiments showed that TO and DMSO stabilized the triplex structures without changing their conformation. In summary, these experiments show that all of the test oligonucleotides formed the desired structure and were sufficiently stable (Tm > 25 °C) to be used in the FID assay.

The affinity of TO for the oligonucleotides in the structure array was determined by fluorimetric titrations carried out under the optimized buffer conditions (76). The results are summarized in Figure 1 and Table S3. The corresponding binding isotherms and emission spectra are in Figures S4-15, panels H and I. The degree of fluorescence enhancement of TO binding depended on the DNA structure (Table S3). In summary, the titrations showed that TO binds with similar affinity (Ka∼9×10^5^-6×10^6^ M^-1^) to all of the oligonucleotide structures with the exceptions of i-motifs 7 and 10 and RNA duplexes 11 and 12, where lower affinities (Ka< 6×10^5^ M^-1^) were observed. Because of their low TO affinity, these structures were omitted from the test panel. A simple 1:1 DNA:TO model provided adequate fits to all of our titration data for the protocol used, except for c-Myc G4 where a secondary binding process was observed at high c-Myc:TO ratios. This weak apparent binding may result from TO-mediated quadruplex dimerization as recently reported for some viral G4s (77). Since the contribution of the low affinity binding was negligible under the conditions of the FID assay, this low-affinity process was ignored in evaluating equilibrium constant for TO:c-Myc G4 interaction.

**Figure 1.**
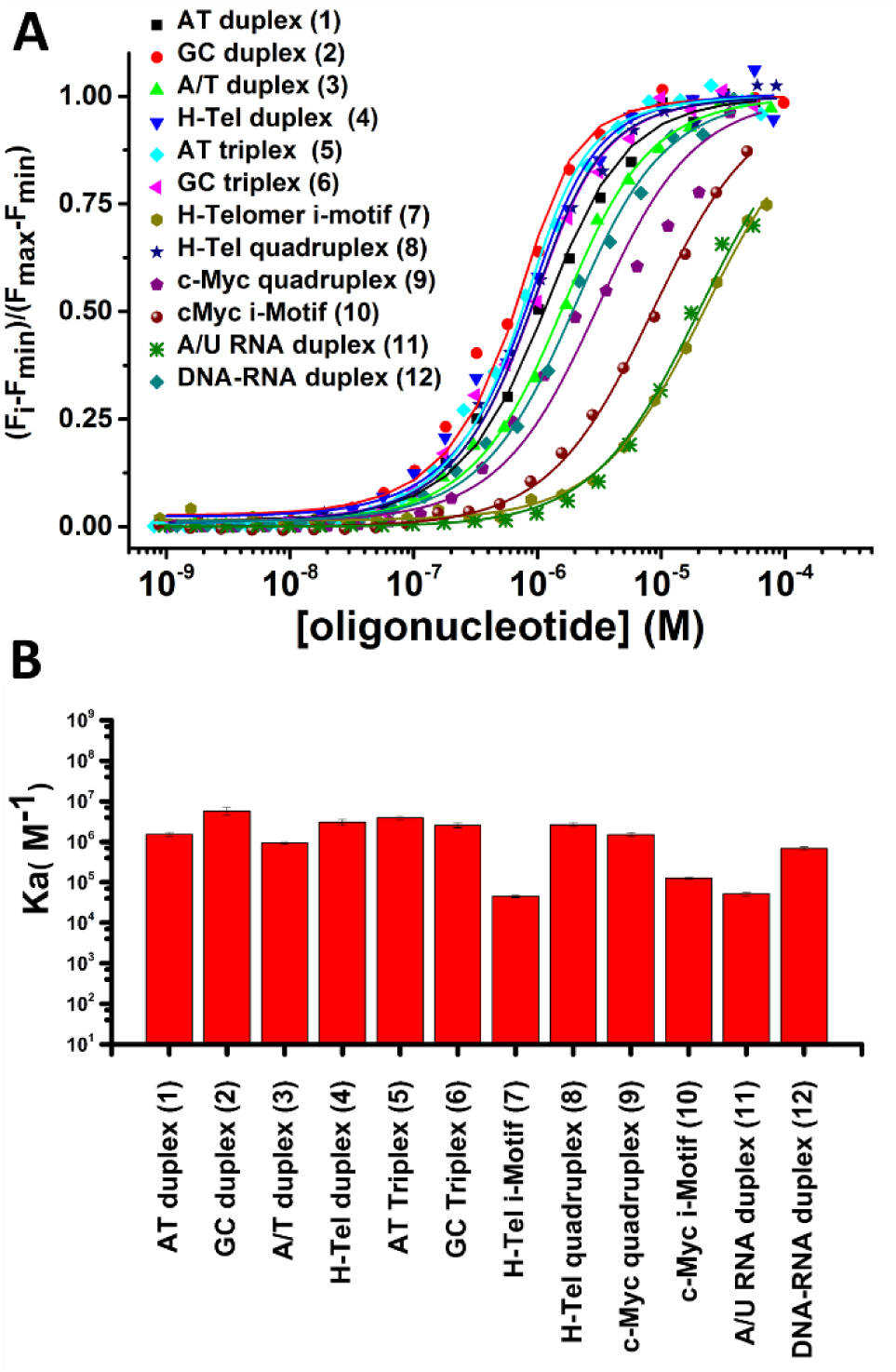
(A) Saturation curves of the fluorescence titrations of TO with different oligonucleotides. (B) Bar graph showing binding constants of TO for different DNA structures used in this study

### FID Assay Design

Assay conditions were optimized to give high sensitivity, a high signal-to-noise ratio, and to minimize the effects of different binding stoichiometries. Corrections were applied to account for any interference from intrinsic ligand fluorescence or from inner filter effects. To obtain the maximum sensitivity, the gain of the microplate reader was set to maximize the fluorescence of AT triplex 5 which exhibited the largest change in emission intensity. The reagent concentrations of 1μM TO and 2 μM nucleic acid were selected because these concentrations gave ∼75 % saturation (Figure 1), a point in the binding isotherm sensitive to competitive displacement and which minimizes potential problems from the binding of multiple probe molecules to the structure (26,56). The lower oligonucleotide/TO ratio used in this work differs from previous assays where excess TO was used (27,28,49,55,56). The lower ratio ensures that only ligands with high affinity will be detected. Figure S19 shows a simulation of the expected TO displacement as a function of ligand binding affinity. This simulation uses the exact concentrations in our assay and the experimentally determined TO binding constants for each structure. The simulations show that under the conditions of our assay the onset of TO displacement requires a ligand affinity of >10^5^ M-1, and complete displacement requires a ligand affinity of >10^8^ M-1.

The effect of test compound concentration was also examined. The results show that good discrimination between oligonucleotides in the library was obtained at a compound concentration of 5 μM (Figure S20).

We also assessed the inner filter effect of fluorescent ligands ethidium bromide, doxorubicin, NMM580, TmPyP2 and TmPyP4 and a non-fluorescent ligand, netropsin, which was selected as a negative control because its absorption spectrum does not overlap with the absorption or emission spectra of TO. The results show that the absorbance of these compounds at 5 μM is <0.06 at the wavelengths of TO excitation (500 nm) and emission (527 nm) (Figure S21-22). Therefore, the inner filter effect is negligible. For instance, 5 μM doxorubicin or TmPyP2 have absorbances at 500 nm and 528 nm that are similar (Figure S21) but their %FID profiles for individual duplex structures differs (Figure S 20 panel C and E). This shows that the difference in binding selectivity for these structures results from different affinities rather than an inner filter effect. In addition, the importance of assessing the spectroscopic properties of potential ligands for possible interference with the FID assay, is illustrated by a %FID > 100% for TmPyP4 (Figure S20 F). This apparent anomaly most likely results from fluorescence of the ligand at the emission wavelength of TO (Figure S22).

### FID assay: reference library

We validated the FID assay for nucleic acid structural selectivity with a reference library of 30 nucleic acid ligands tested against eight oligonucleotide structures. The ligand library contains various intercalators (*e.g*. ethidium bromide), minor groove binders (e.g. netropsin), triplex binders (*e.g.* coralyne) and G4 ligands (*e.g*. NMM) (Table S5, Figure S23). In addition, a variety of other chemical structures including heterocycles, carbohydrates, and peptides, each with different physical, chemical, spectroscopic and binding properties, were in the reference library.

The results of the FID assay of the 30 ligands binding to eight nucleic acid sequences and structures (4 duplex, 2 triplex and 2 quadruplex) is presented as a bar graph in Figure 2. The data report the results of 240 individual binding interactions. Positive and a few negative values of %FID were observed. Positive values can be explained by a fluorescence decrease resulting from displacement of TO from the nucleic acid by the competitor ligand. In these cases, %FID is directly related to the affinity of a compound for a given nucleic acid structure. The higher the %FID, the higher affinity of the compound for a nucleic acid structure. For example, netropsin showed 84 %FID for AT duplex 1 compared to 6% for GC duplex 2. This is consistent with the known selectivity of netropsin for AT over GC duplexes (23,25).

**Figure 2.**
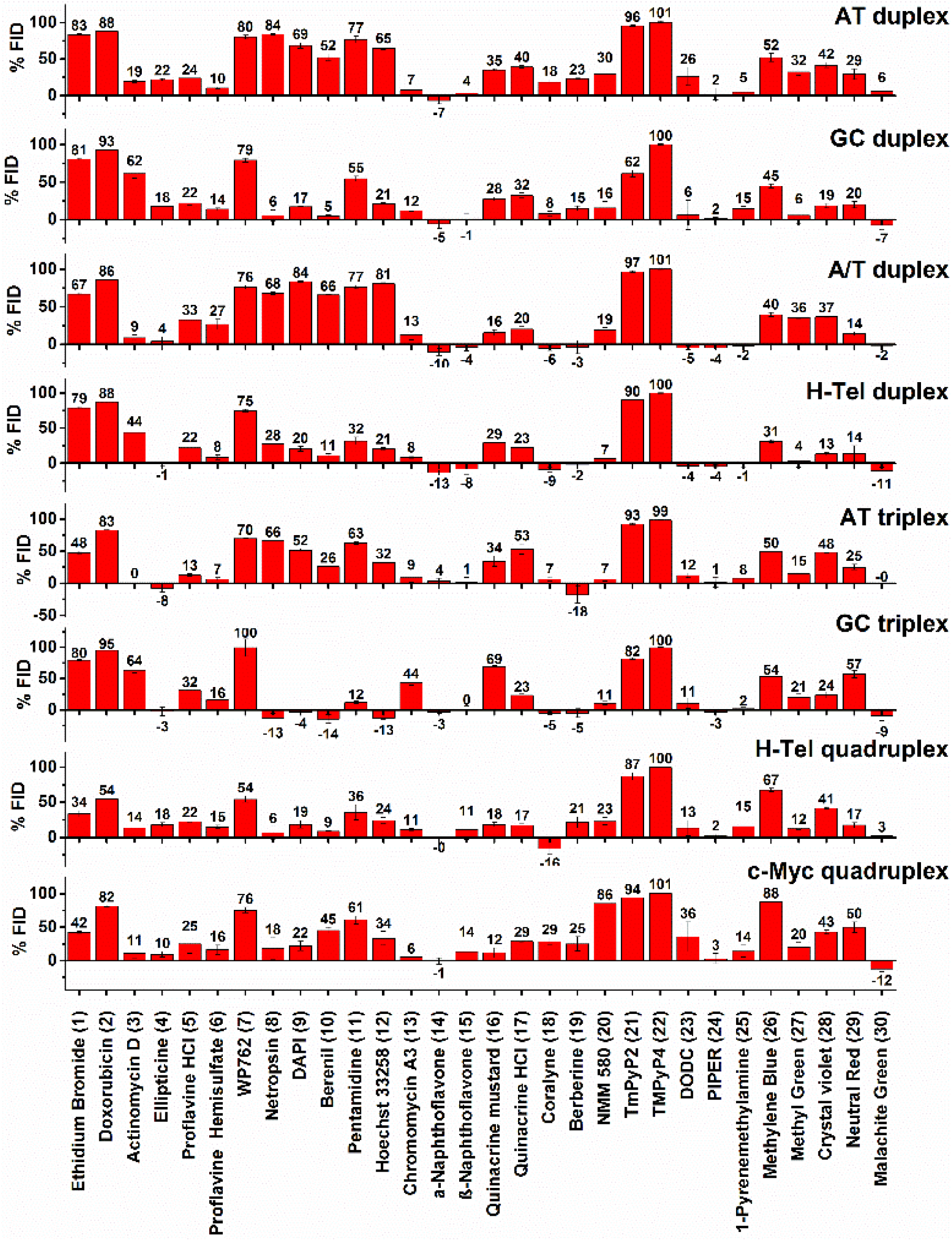
Results of FID assay performed with 30 nucleic acid binders and 8 oligonucleotide structures. Each bar represents the percentage of TO displacement (%FID).

Negatives values of %FID are more difficult to rationalize (54). A number of factors might account for TO fluorescence enhancement leading to a negative %FID: a) interaction between TO and the test compound; b) increased binding affinity of TO induced by the test compound; c) similar photo-physical behavior of the compound with TO. To further investigate the negative %FID observed with *α*-naphtoflavone, coralyne and berberine interacting with the A/T duplex, the AT triplex and H-tel G4 structures, we determined absorbance, excitation and emission spectra of TO in the presence and absence of ligand under the same experimental conditions used for the FID (Figures S 24-28). Both α-naphthoflavone (Figure S24) and coralyne (Figures S25-26) enhanced TO fluorescence both in the presence and absence of oligonucleotide. This accounts for the observed negative %FID and suggests an interaction between these compounds and TO. For berberine (Figure S27-28), significant differences were observed in the fluorescence spectra when the experiment was conducted in a quartz cuvette compared to a 96-well plate. The %FID observed for berberine using a quartz cuvette was -1%, in contrast to -15 % observed when a 96-well plate was used. This suggests non-specific binding of berberine to the plastic plate.

### Statistical data analysis: principal component and cluster analysis

Traditional methods of reporting FID HTS data such as bar graphs are cumbersome for detailed specificity analysis since it is difficult to grasp global trends. A major goal of the present study was to develop an efficient, quantitative approach for more effectively evaluating ligand-nucleic acid interactions. The methods of principle component analysis (PCA) and hierarchal cluster analysis (HCA) provide the tools for such al method.

PCA is a multivariate statistical method to reduce the dimensionality of data without significant loss of information using derived variables (principal components) (78-81). This is a non-supervised classification method that performs unbiased clustering of the data. PCA is particularly useful for finding hidden patterns and aiding in interpretation of high dimension, complex data sets (82). PCA projects the multi-dimensional data onto a new lower dimension subspace. The axes in this new subspace, known as principal components (PCs), are linear combinations of the original variables with their origin located at the center of the multi-dimensional data. The PCs are independent (orthogonal) to each other and retain as much of the information in the original variables as possible. The first principal component (PC1) corresponds to the axis that contains the greatest variation of the data. The second principal component (PC2) is orthogonal to the first and oriented in the direction of the second greatest amount of variation of the data, and so on.

In addition to PCA, we used HCA as an unsupervised classification method to find clusters of compounds. HCA calculates the distance between observations in multidimensional space and forms clusters of individuals based on the similarity of their variables.

The results of PCA and HCA performed on the FID data of our reference library of 30 nucleic acids binders and 8 oligonucleotide structures are shown as biplot graphs, PC1-PC2 (Figure 3, left panel) and PC2-PC3 (Figure 3, right panel) (83). The variables (oligonucleotides) are represented by arrows and the individuals (compounds) by points colored according to their cluster. The results of HCA performed on the PCs, after consolidation by an additional partitional clustering with a K-means algorithm (84), are shown as dendrograms in Figures S29 and S30. The horizontal axis represents the compounds and the vertical axis measures the distance or similarity between individuals in the clusters. The more similar the affinity and selectivity of the individual ligands for the different DNA structures, the lower are the nodes in the tree. The levels at which the hierarchical tree was cut to initialize the k-means algorithm are represented by rectangles.

**Figure 3.**
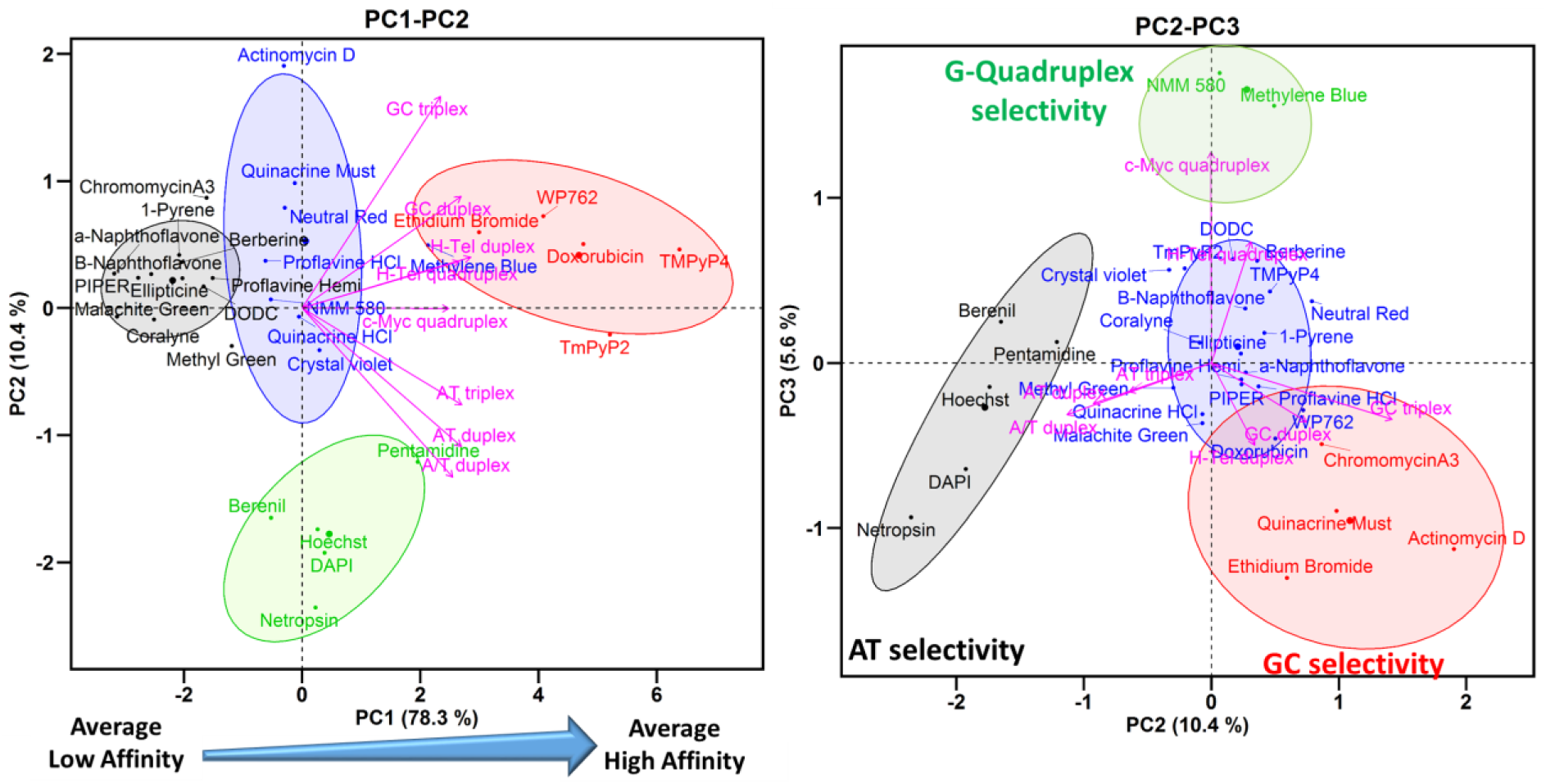
PCA and HCA of FID results for 30 nucleic acid binders and 8 oligonucleotide structures. Left: biplot of PC1 and PC2; right: biplot of PC2 and PC3. The center of each cluster is represented by larger point size.

### Interpretation of the PC1-PC2 biplot

Four well-separated clusters are observed in the biplot of PC1-PC2 (Figure 3, left), which explains 89% of the variance of the data. Cluster 1 (black) includes the compounds that exhibit low or null capacity for displacement of TO (low %FID) from all the nucleic acid structures. Cluster 2 (blue) and cluster 3 (green) contain compounds situated close to the origin of PC1 and which showed medium affinity. Cluster 3 (green) contains minor groove binders pentamidine, netropsin, berenil, DAPI, Hoechst that have a selectivity for AT sequences (23,25). Cluster 4 (red) contains DNA intercalators ethidium bromide, doxorubicin, WP762 and porphyrins TmPy4, TmPy2 that exhibit high values of %FID for almost all of the DNA structures.

Analysis of the loading factors (the magenta arrows) reveals that all of the oligonucleotides have a positive correlation and similar contribution to PC1 as indicated by the length and direction of projection of the variables on to PC1 (Figure 3, left). The biplot PC1-PC2 shows that oligonucleotides with similar AT or GC base composition cluster together along PC2. Low correlation between AT-rich (AT triplex, AT duplex and A/T duplex) and GC-rich (GC triplex and GC duplex) oligonucleotides is observed along PC2. H-Tel duplex, H-Tel G4 and c-Myc G4, with similar AT and GC content (∼50 % AT), are situated between the AT-rich and GC-rich oligonucleotides. PC1 therefore can be interpreted in terms of global affinity for different DNA structures and PC2 can be interpreted as showing different ligand affinities for AT- or GC-rich oligonucleotides.

The projection of a compound onto a particular oligonucleotide vector is interpreted as its relative affinity for a particular DNA structure. For instance, the projections of the compounds WP762, methyl blue, NMM580, methyl green onto the GC duplex vector within the PC1-PC2 biplot (Figure 3, left), shows that the order of affinity for GC duplex (2) is approximately WP762 > methyl blue > NMM580 > methyl green, which agrees with the affinity ranking obtained from the %FID.

In summary, the biplot PC1-PC2 allows easy visualization and classification of compounds based on their affinity for the DNA structures. PC1 separates the compounds based on their global affinity and PC2 separates them based on their AT or GC affinity. In this way, compounds situated at the left, center and right correspond to compounds with high, medium or low affinity for the different DNA structures.

### Interpretation of the PC2-PC3 biplot

The PC2-PC3 biplot (Figure 3, right) projection provides visualization of the selectivity of a compound for a particular DNA structure. Three groups of variables hidden in the PC1-PC2 biplot are now observed in the PC2-PC3 biplot (Figure 3, right). Each group of variables points in different directions with an angle between them of ∼120°. The first group contains AT-rich structures (AT duplex, A/T duplex and AT triplex) and points to the bottom left corner of the PC2-PC3 biplot. The second group contains GC-rich structures (GC duplex, GC triplex and H-Tel duplex) and points to the bottom right corner. The third group contains only the G4 structures H-Tel and c-Myc and points in a positive direction of PC3. This orientation of the variables reflects differences in the structures of these oligonucleotides. Therefore, information about the selectivity for a particular DNA structure can be obtained from the location of the compounds in the PC2-PC3 biplot.

The biplot PC2-PC3 shows four well-separated clusters of compounds. Cluster 1 (black) contains the minor groove binders pentamidine, netropsin, berenil, DAPI and Hoechst that are selective for AT in duplexes and triplexes (25). Cluster 2 (red) contains the DNA intercalators ethidium bromide, actinomycin D, doxorubicin, WP762 and also the minor groove ligand chromomycin A3 that prefers GC base pairs (85). These compounds have higher affinity for GC-rich DNA structures (25). Cluster 3 (green) contains NMM580 and methylene blue that are selective for G4s (21,25,86,87). Cluster 4 (blue) includes the remaining compounds that have low selectivity. To summarize, the PC2-PC3 biplot shows a higher level of discrimination between different DNA structures. PC2 separates compounds based on AT or GC base composition and PC3 discriminates based on G4 binding.

### Application to the NCI diversity set III library

In order to demonstrate the utility of this FID assay as a secondary screen in the context of a drug discovery platform, we used the NCI diversity set III library that contains 1598 compounds as a test. As a primary screen of the library, differential scanning fluorometry was used to identify compounds that bound to a FRET-labeled G4 structure, a human telomere sequence 5’AGGG(TTAGGG)_3_ in the hybrid form. Figure S31 shows the distribution of T_m_ shifts obtained, with several compounds with ΔT_m_ > 10 degrees, indicative of tight binding. We selected 20 of these with T_m_ shifts ranging from 6 to 31 degrees (Figure S31-32) for secondary screening using HT-FID, based on their apparent affinity and compound availability. The results of this screen are shown in Figure 4.

**Figure 4.**
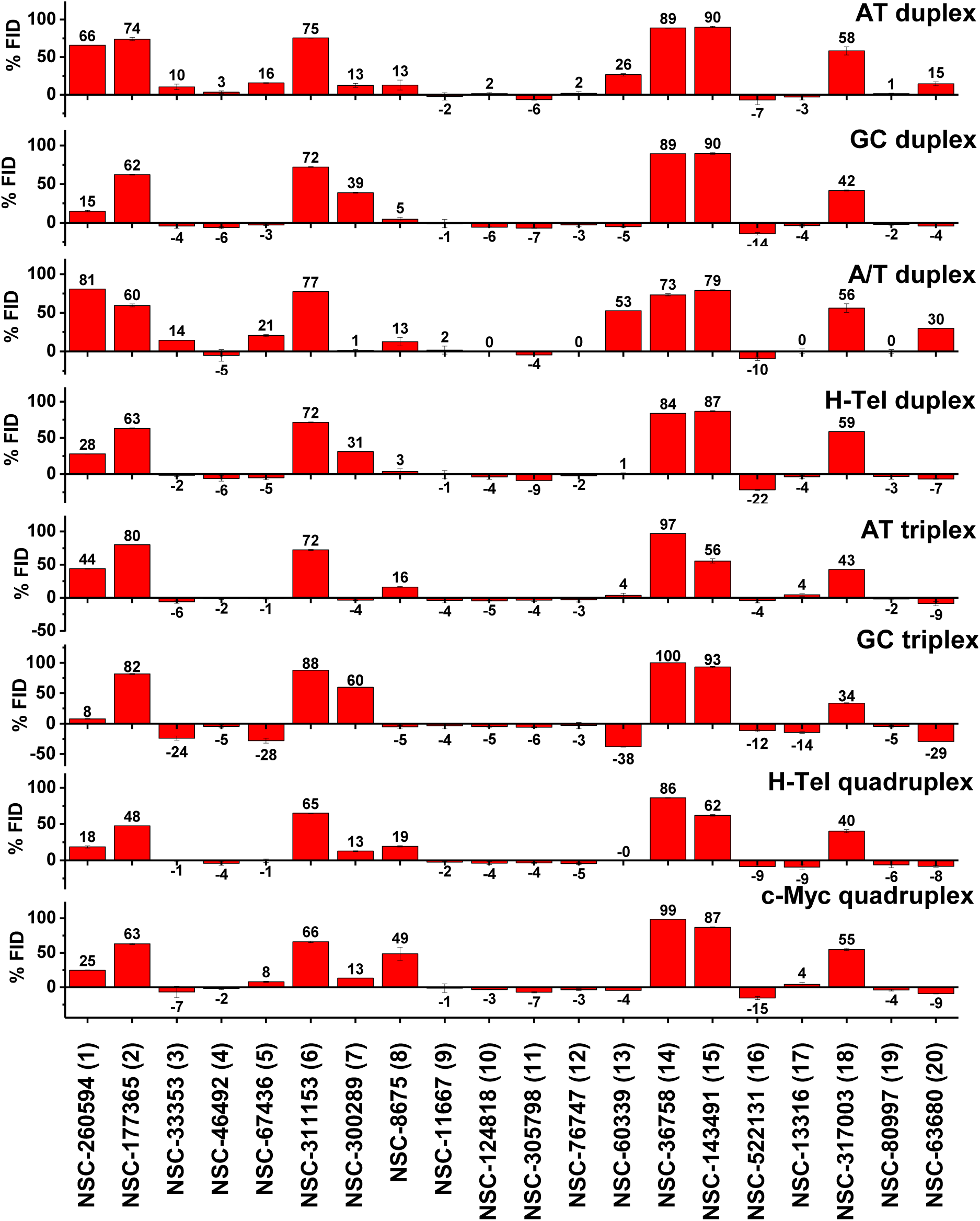
Results of the FID assay of 31 compounds of the NCI diversity set III library and 8 oligonucleotide structures. Each bar represents the % FID of a given compound for an oligonucleotide sequence.

To our surprise, not all of the 20 compounds that registered as hits in the thermal shift assay were able to displace TO from the unlabeled G4 hybrid structure used in our FID assay, even though the identical sequence was used in both assays. No correlation between the magnitude of ΔT_m_ and the %FID recorded (Figure S33) was observed. To understand this result, several differences between the FID and thermal shift assays need to be emphasized. First, the thermal shift assay requires a FRET labeled G4 structure, which might influence binding, while the FID assay does not. Second, the thermal shift assay uses a large molar excess of ligand over G4 (250:1), while a much lower molar ratio (5:1), is used in the FID assay, perhaps resulting in differing extents of ligand binding. Third, the FID assay is by design a competition binding assay and the apparent ligand affinity is relative to the affinity of the TO probe and is reduced from the true affinity. Recall that, as discussed above, a ligand binding constant of >10^5^ M^-1^ is required to register TO displacement in the FID assay.

Even with these considerations, the lack of correlation between the thermal shift and FID assays remains puzzling. Additional explanations for the differences are needed. First, it is possible that the ligand may bind to a different site than TO and instead of displacement, a ternary (ligand-G4-TO) complex may form. For example, if TO is bound by an “end pasting” mode and stacked on a terminal G-quartet, the ligand might bind within a groove and not perturb the bound TO. Second, it is possible that the FRET labels in the thermal shift assay might somehow form a binding pocket to facilitate ligand binding, and since they are not present in the FID assay, no ligand binding site would be available.

We conclude that a positive FID confirms and validates ligand binding registered by the primary thermal shift assay, but that a valid initial hit might be invisible in the FID assay. Since the intent of our FID assay is as a secondary screen to assess binding selectivity, we focused further analysis on those compounds that showed appreciable FID (>10%) in their interaction with the human telomere hybrid G4. Five of these eight compounds with an FID response had ΔT_m >_ 25°C in the primary screen.

The most striking result to emerge from inspection of Figure 4 is that only 1 of the 8 G4 hybrid binders exhibited any specificity. Compounds 2,6,14,15 and 18 (Figure 4, Figure S32) bind to all structures in the FID assay. Compound 1 has a selectivity for AT-oligonucleotides and a shape similar to the classical minor groove binders. Compound 7 is interesting because it shows a clear preference for GC-oligonucleotides, with significant binding to the GC triplex structure and GC-rich duplexes in addition to binding to the two quadruplex structures. Compound 8 emerges as the most selective compound for G4 structures. Interestingly, although it was selected for binding to an antiparallel hybrid G4 structure, it clearly binds preferentially to the c-myc parallel G4 structure. It shows some binding to AT-rich triplex and duplex structures, but not to GC-rich duplex or triplex structures. The key point is that the FID assay fulfilled its intended purpose as a secondary screen for binding specificity. While it is disappointing that most hits from the primary thermal shift assay seem to lack specificity, it is essential to find that out sooner rather than later to avoid further development of less than ideal compounds for targeting G4 structures. Perhaps such lack of specificity is a contributing reason for the unsuccessful discovery of G4 targeted drugs so far.

A PCA analysis provides further insight into the selectivity of the eight compounds, with the results shown in Figure 5 (Figure S34-35). In this analysis, the FID data of Figure 4 for the 8 compounds showing TO displacement for the hTel G4 was combined with the FID data for the 30 reference compounds. The interesting results that emerge are that compound 8 clusters with known quadruplex binders, compound 1 clusters with known groove binders, and compound 7 clusters with known GC-specific intercalators. The remaining five compounds have little or no selectivity. These results illustrate the utility of the FID assay and PCA analysis for defining specificity and possible binding modes. Inspection of the structure of compound 8 shows that it is similar to crystal violet that is in our set of reference compounds, but seems to have improved selectivity for G4 binding.

**Figure 5.**
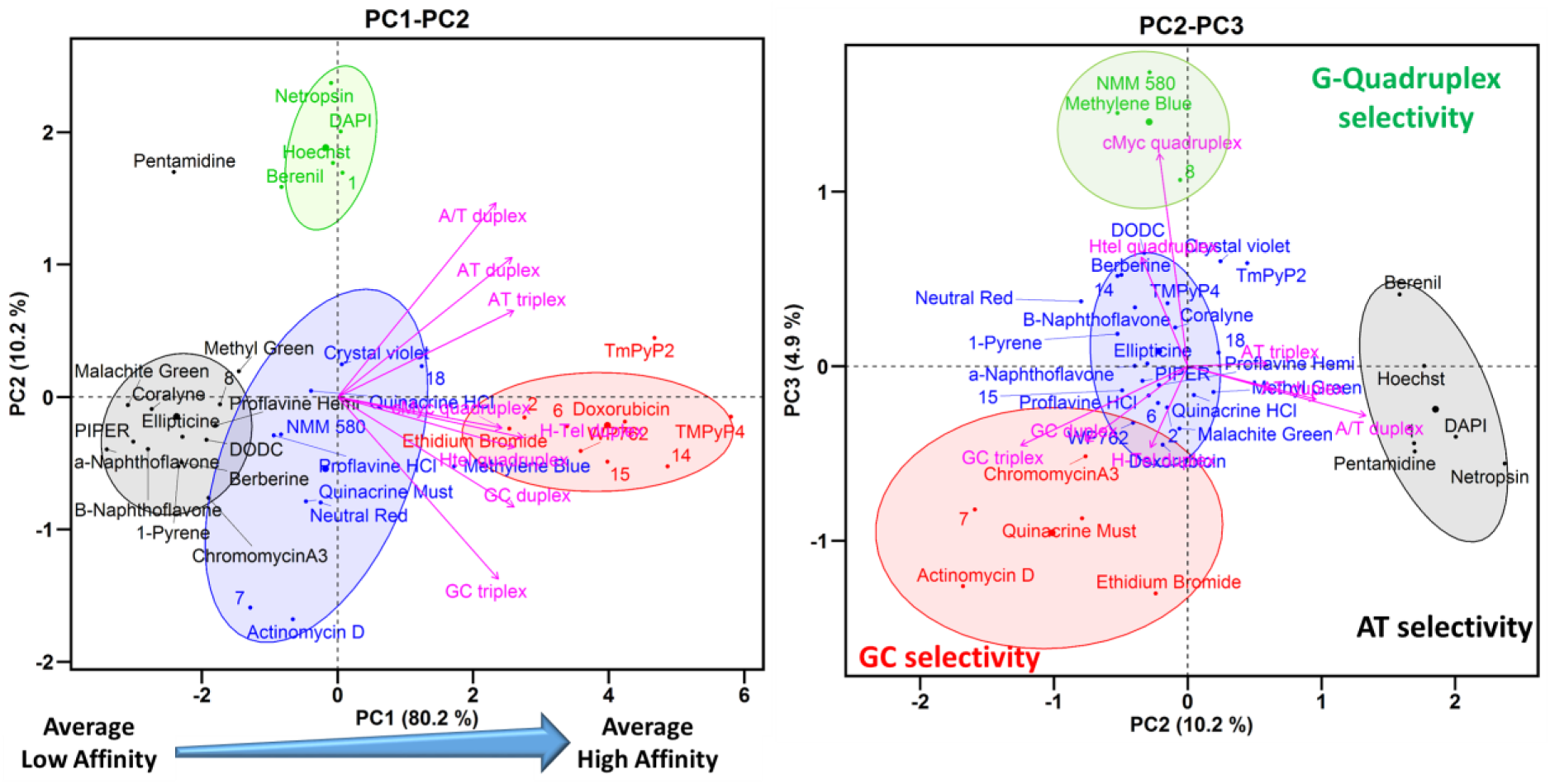
PCA and HCA of FID results for the combination of 30 nucleic acids binders and the 8 compounds with the highest thermal shift (ΔTm) and % FID for H-Tel quadruplex from the NCI diversity set III library with 8 oligonucleotide structures. Left: biplot of PC1 and PC2. Right: biplot of PC2 and PC3. The center of each cluster is represented by larger point size.

## SUPPLEMENTARY DATA

Supplementary Data are available at NAR Online.

